# An artificial cell containing cyanobacteria for endosymbiosis mimicking

**DOI:** 10.1101/2021.04.08.438728

**Authors:** Boyu Yang, Shubin Li, Wei Mu, Zhao Wang, Xiaojun Han

**Affiliations:** State Key Laboratory of Urban Water Resource and Environment, School of Chemistry and Chemical Engineering, Harbin Institute of Technology, 92 West DaZhi Street, Harbin, 150001, China

## Abstract

The bottom-up constructed artificial cells help to understand the cell working mechanism and provide the evolution clues for organisms. Cyanobacteria are believed to be the ancestors of chloroplasts according to endosymbiosis theory. Herein we demonstrate an artificial cell containing cyanobacteria to mimic endosymbiosis phenomenon. The cyanobacteria sustainably produce glucose molecules by converting light energy into chemical energy. Two downstream “metabolic” pathways starting from glucose molecules are investigated. One involves enzyme cascade reaction to produce H_2_O_2_ (assisted by glucose oxidase) first, followed by converting Amplex red to resorufin (assisted by horseradish peroxidase). The more biological one involves nicotinamide adenine dinucleotide (NADH) production in the presence of NAD^+^ and glucose dehydrogenase. Further, NADH molecules are oxidized into NAD^+^ by pyruvate catalyzed by lactate dehydrogenase, meanwhile, lactate is obtained. Therefore, the sustainable cascade cycling of NADH/NAD^+^ is built. The artificial cells built here simulate the endosymbiosis phenomenon, meanwhile pave the way for investigating more complicated sustainable energy supplied metabolism inside artificial cells.

## Introduction

The origin of eukaryotic cells is a fatal puzzle of evolution, which shows great significance to elucidate the relationship among the internal structures of eukaryotic cells^1,2^. There are two representative theories of eukaryotic cell origin including “classic theory” and “endosymbiosis theory”, among which the latter is far more widely recognized in academia^3,4^. The “endosymbiosis theory” considers that chloroplasts originate from cyanobacteria^5,6^. The most studies on endosymbiosis focus on genome analysis to find the connection between chloroplast and cyanobacteria^7-9^. To the best of our knowledge, there is no report on mimicking endosymbiosis in an artificial cell system.

Artificial cells are defined as the bottom-up built structures which mimic one or more cell characteristics (membrane, metabolism, reproduction, etc.)^10,11^. Those structures help to deep understand the working mechanism of cells. The metabolism mimicking in artificial cells initially mainly relied on enzymatic cascade reactions in the “cytoplasm”^12,13^ or between two protoorganelles^14,15^. The sustainable energy supply for metabolism attracted more and more attentions^16,17^. The protoorganelles with energy harvesting capability to produce adenosine triphosphate (ATP) were designed to mimic chloroplasts^18^. The long-term sustainable energy supply to simulate the process of metabolism in synthetic cells is still a big challenge.

Targeting the issue of sustainable energy supply in artificial cells, as well as inspired by endosymbiosis theory, herein, an artificial cell containing cyanobacteria was constructed, where cyanobacteria were “chloroplasts” for sustainable energy supply. Two metabolism pathways were investigated as shown in Figure 1, both of which started from the glucose produced by cyanobacteria. One was the cascade enzyme reactions involving glucose oxidase (GOx) and horseradish peroxidase (HRP). The glucose molecule was oxidized by GOx to produce H_2_O_2_ (equation 1 in Figure 1b), which further oxidize Amplex red to resorufin in the presence of HRP (equation 2 in Figure 1b)^19^. The other pathway was to synthesize the “energy currency” NADH from NAD^+^ and glucose catalyzed by glucose dehydrogenase (GDH) (equation 3 in Figure 1b). With the energy supply of NADH, pyruvate was transformed into lactate catalyzed by lactate dehydrogenase (LDH) (equation 4 in Figure 1b). The produced NAD^+^ reentered the previous reaction (equation 3) to form NADH recycle. The artificial cell provides an advanced cell model for mimicking endosymbiosis with complicated intracellular energy supply for cell metabolism.

**Figure 1.**
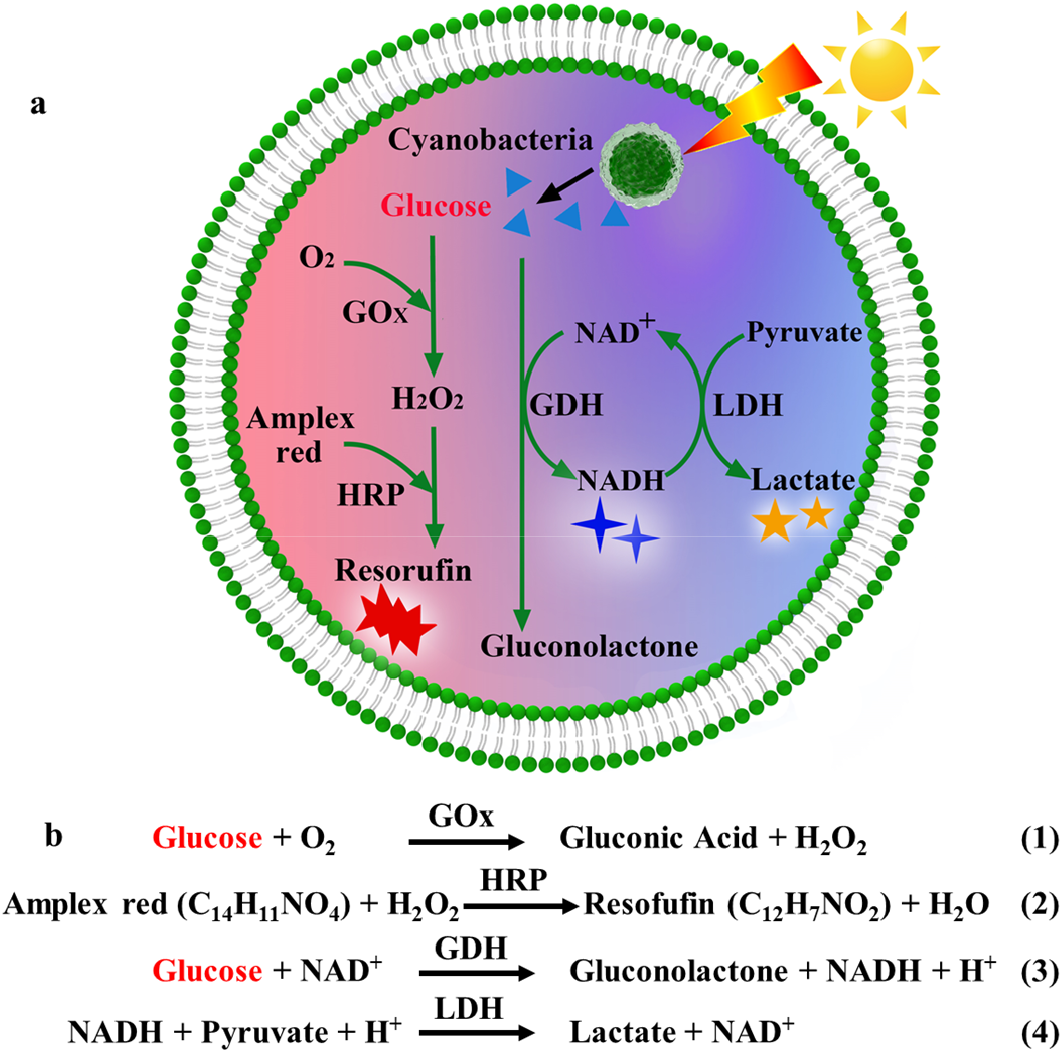
Schematic illustration of endosymbiosis mimicking artificial system. (a) Scheme of cyanobacteria containing artificial cell for endosymbiosis mimicking. (b) Chemical reaction equations involved in metabolism mimicking pathways.

## Results

### Preparation of artificial cells containing cyanobacteria

Cyanobacteria can be directly observed by the fluorescence microscope due to intracellular chlorophyll (Figure S1a). The diameter of cyanobacteria is 3.55 ± 0.39 μm from bright field microscopy images (Figure S1b, c). Their zeta potential is -26.9 ± 1.2 mV. Cyanobacteria were encapsulated into giant unilamellar vesicles (GUVs) to form artificial cells using the electroformation method (Figure S2). The cyanobacteria outside the artificial cells were removed by filtering the solution through a polycarbonate membrane with the pore size of 10 μm in diameter. The number of cyanobacteria inside the artificial cells was roughly controllable by varying the concentration of cyanobacteria in the solution during the electroformation. The encapsulation efficiency is defined as the number of GUVs containing cyanobacteria divided by the total number of GUVs at certain cyanobacteria concentration used for artificial cell preparations. The encapsulation efficiency (Fig. 2a) gradually increased from 3.4 ± 1.4% (7.7 mg/mL) to 41.9 ± 14.2% (306.1 mg/mL) with the increase of cyanobacteria concentration. At the same concentration, the artificial cells contained different number of cyanobacteria. We classified the artificial cells into two groups according to the number of cyanobacteria inside the artificial cells to be less than 5 or not. Below the concentration of 30.6 mg/mL, it is less chance to obtain the artificial cells containing more than 5 cyanobacteria (Figure 2b). The fraction of those (with more than 5 cyanobacteria) increased to 44.4% at high concentration of 306.1 mg/mL. The typical fluorescence images of artificial cells containing one cyanobacterium and more than 5 cyanobacteria were presented in Figure 2c and d respectively. Those containing two, three and four cyanobacteria were also obtained (Figure S3). The confocal cross-section images in Figure 2e confirmed the cyanobacteria encapsulated inside the artificial cell rather than on the top or bottom of the artificial cell. Movie S1 showed the cyanobacteria moved up and down freely inside the artificial cell, which also confirmed the encapsulation of cyanobacteria. At the concentration of 22.9 mg/mL, a large fraction (>79.1%) artificial cell contained one cyanobacterium. Therefore, in the following experiments, we choose cyanobacteria concentration to be 22.9 mg/mL to obtain the artificial cells.

**Figure 2.**
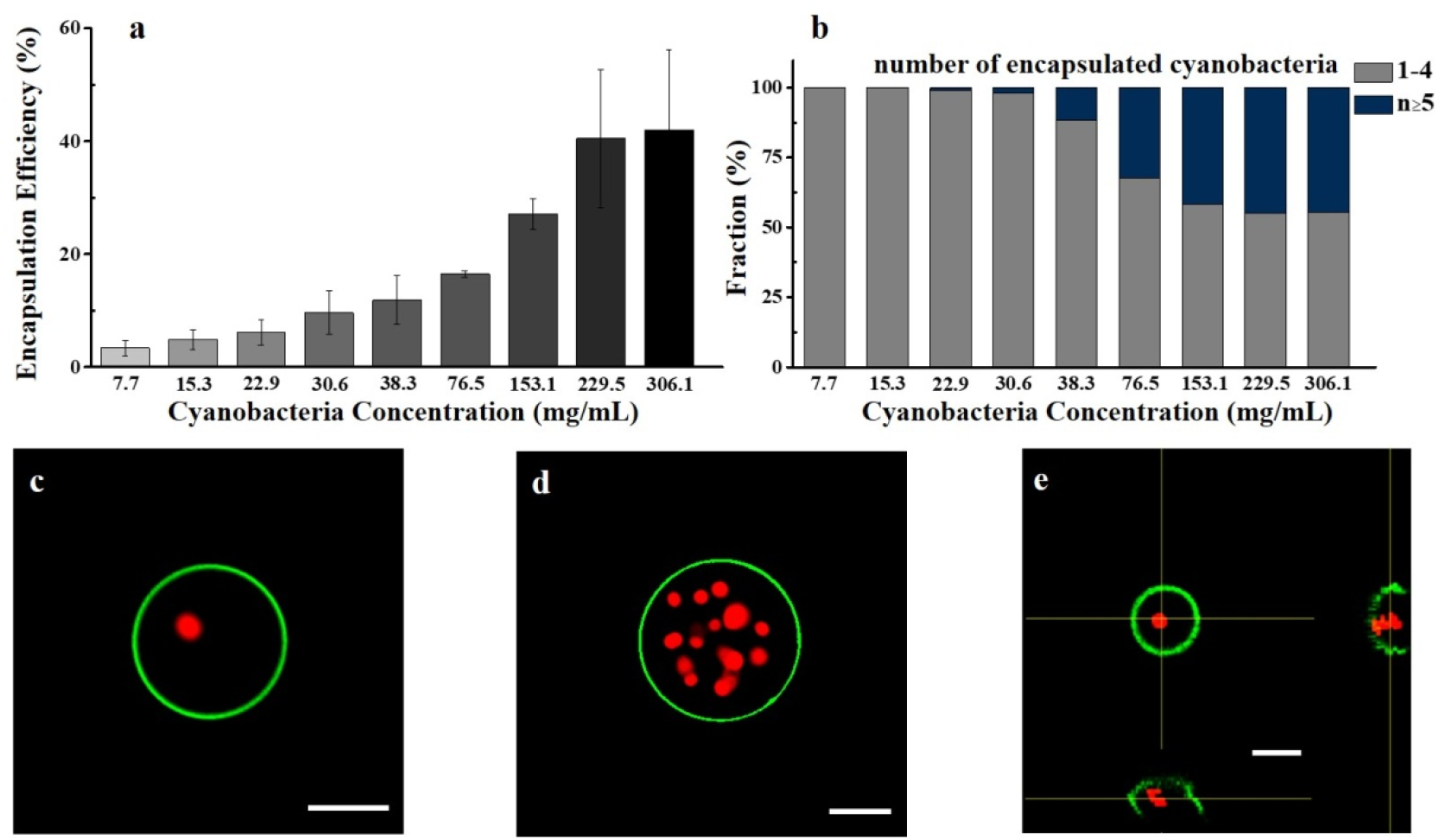
Encapsulation of cyanobacteria inside GUVs. (a) Encapsulation efficiency as a function of cyanobacteria concentration. The encapsulation efficiency was 3.4 ± 1.4%, 4.9 ± 1.7%, 6.1 ± 2.2%, 9.6 ± 3.8%, 11.9 ± 4.3%, and 16.5 ± 0.6%, 27.1 ± 2.8%, 40.5 ± 12.2% and 41.9 ± 14.2% with the cyanobacteria concentration of 7.7, 15.3, 22.9, 30.6, 38.3, 76.5 153.1, 229.5 and 306.1 mg/mL, respectively. Data are presented as mean values ± SD. Error bars indicated standard deviations (n = 3). (b) The fraction of artificial cells containing less than 5 cyanobacteria and more than 5 cyanobacteria at different cyanobacteria concentrations. The fraction of artificial cells containing less than 5 cyanobacteria was 100%, 100%, 99.1%, 97.9%, 88.5%, 67.5%, 58.3%, 55.5% and 55.0% with the cyanobacteria concentration of 7.7, 15.3, 22.9, 30.6, 38.3, 75.6, 153.1, 229.5 and 306.1 mg/mL, respectively. At least 100 GUVs were counted to obtain each data. Representative fluorescence microscopy images of the artificial cells containing a single (c) and multiple (d) cyanobacteria. (e) The confocal image of the artificial cell containing one cyanobacterium with two cross-section images. All scale bars were 20 μm. The artificial cells were labeled with NBD-PE in the lipid bilayer (green fluorescence).

### Cyanobacteria as an “organelle” for continuous energy supply

Cyanobacteria are considered to be the ancestor of chloroplasts according to endosymbiosis hypothesis. In our prepared artificial cells, cyanobacteria were used as organelles to mimic chloroplasts, which converted light energy into chemical energy. One of the metabolism products of cyanobacteria upon light illumination is glucose. The glucose kit was used in the test tube to monitor the real-time production glucose when the cyanobacteria underwent photosynthesis. From 0 to 24 h, the concentration of photosynthesized glucose continuously increased against time, as shown in Figure 3c. After 24 h illumination by daylight lamp, the photosynthesized concentration of glucose produced by cyanobacteria (22.95 mg/mL) was 4.98 ± 0.21 μg/mL. The solution used for culturing cyanobacterial cells was a mixture of BG-11 medium and sucrose solution (8/2 = v/v), which was the same solution inside the artificial cells. Therefore, it is suggested the glucose can be produced inside the cyanobacteria-containing artificial cells upon light illumination. To confirm this, the following experiments were carried out.

**Figure 3.**
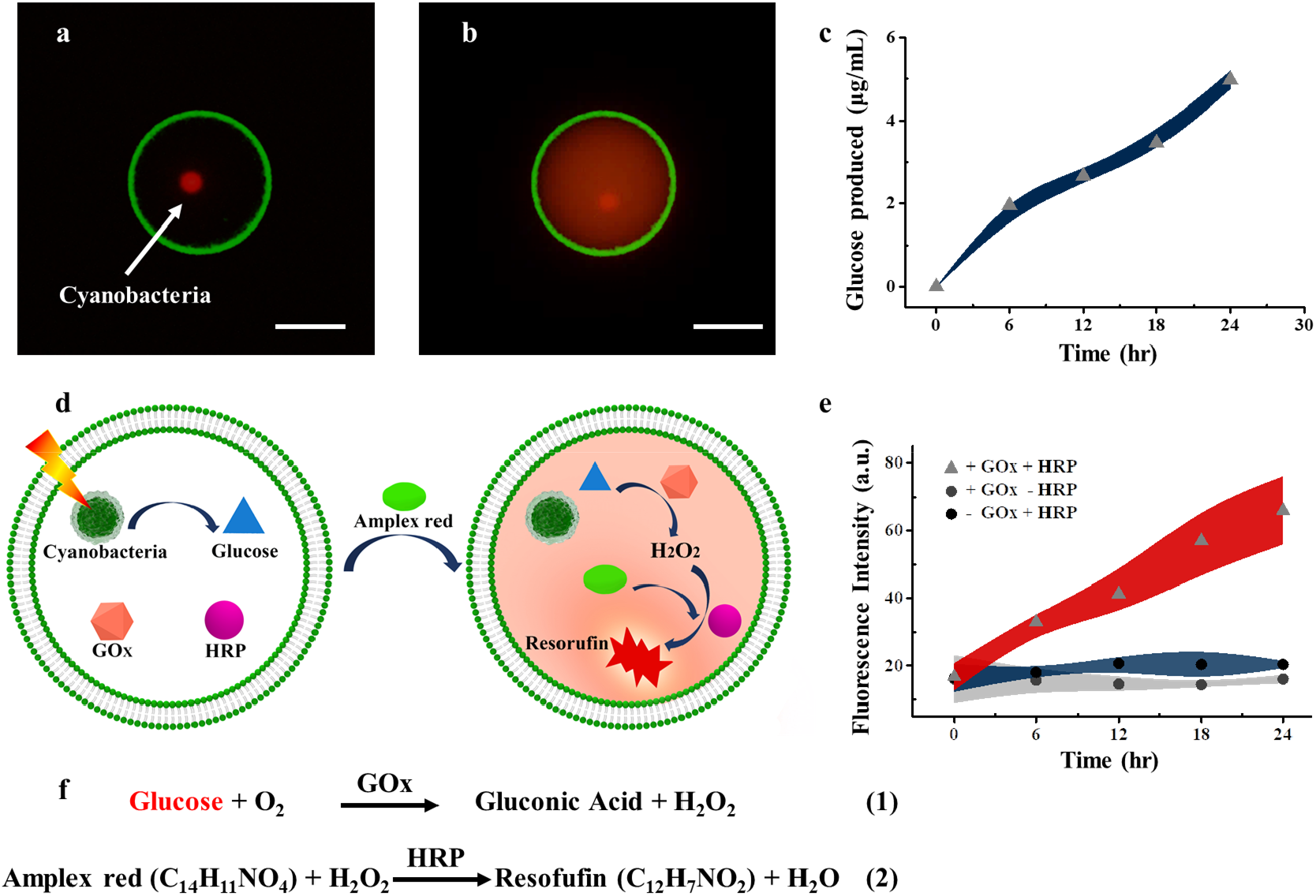
Chemical communications between cyanobacteria and “cytoplasm” inside the artificial cells. The representative images of artificial cells containing cyanobacteria, GOx and HRP before (a) and after (b) the introduction of Amplex red and 24 hours light illumination. (c) The amount of produced glucose as function of illumination time in the test tube. (d) The corresponding scheme of cascade reactions between the cyanobacteria and “cytoplasm” inside an artificial cell. (e) Fluorescence intensity of produced resorufin inside the artificial cells (containing GOx and HRP, GOx only and HRP only) against time, as well as the control experiments. (f) Chemical reaction equations involved in artificial cells. The scale bars were 20 μm. The colored bands indicate mean ± s.d. of independent experiments.

The chemical communication between cyanobacteria and “cytoplasm” was performed. We produced the artificial cells containing cyanobacteria, glucose oxidase (GOx) and horseradish peroxidase (HRP), as illustrated in Figure 3d (left image). The corresponding fluorescent image was shown in Figure 3a. The red dot is cyanobacterium.

Upon light illumination for 24 hours, Amplex red (10 μM) was introduced to diffuse into the artificial cells. The glucose metabolized by cyanobacteria was oxidized by GOx to H_2_O_2_, which further oxidized Amplex red in the presence of HRP to yield the fluorescent resorufin^20,21^ as illustrated in Figure 3d (right image). The “cytoplasm” (Figure 3b) turned into red color as expected. This result confirmed the production of glucose by cyanobacteria upon light illumination. We also monitored the fluorescence intensity as a function of time to indirectly reflect the amount of glucose produced by the photosynthesis of cyanobacteria inside the artificial cells (Figure 3e). The fluorescence intensity increased with the illumination time, showing the same increasing trend as that in Figure 3c. There were no cascade enzyme reactions observed inside the artificial cells containing no GOx or HRP (Figure 3e). To confirm the cell viability in artificial cells, fluorescein diacetate (FDA) stain assay was implemented. FDA is a non-fluorescent molecule that is hydrolyzed in living cells to produce luminescent fluorescein (green fluorescence). After 36 hours, the cyanobacteria in the artificial cells still alive as shown in Figure S4.

The chemical communication between cyanobacteria and protoorganelle inside the artificial cells was further conducted. The artificial cells containing cyanobacteria and glucose oxidase (GOx) were deformed into vesicle in vesicle structures by hypertonic shock^22^, as illustrated in Figure S5. The inner vesicles containing HRP was considered as the protoorganelle^23^. The representative fluorescence microscopy image and schematic illustration were shown in Figure 4a and Figure 4c (left image), respectively. The red dot in Figure 4a represented cyanobacterium. The artificial cells encapsulating cyanobacteria and protoorganelle were exposed to daylight lamp for 24 h. The glucose from the photosynthetic metabolism of the cyanobacteria generated H_2_O_2_ under the catalysis of GOx in the “cytoplasm” of artificial cell, which went through the protoorganelle membrane together with the added Amplex red^24^. Assisted by the catalysis function of HRP inside the ptotoorganelle, red fluorescent resorufin was produced from Amplex red (Figure 4b, right image of 4c). The chemical communications were realized using this artificial cell between real organelle (cyanobacteria) and protoorganelle.

**Figure 4.**
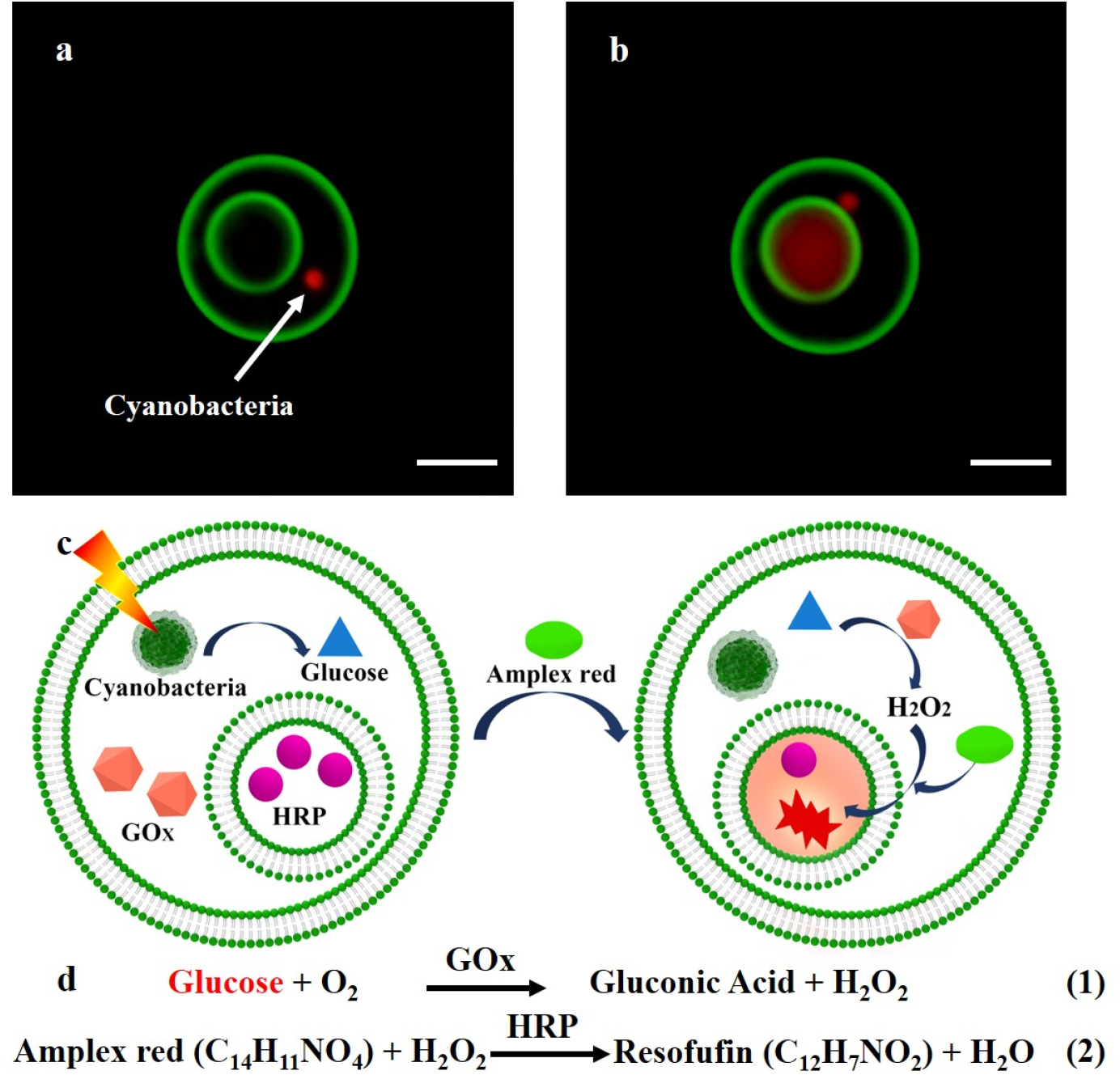
Chemical communications between cyanobacteria and protoorganelle inside the artificial cells. The representative images of artificial cells containing cyanobacteria, GOx and protoorganelle (containing HRP) before (a) and after (b) 24 hours light illumination with the introduction of Amplex red. (c) The corresponding scheme of cascade reactions between the cyanobacteria and protoorganelle inside an artificial cell. (d) Chemical reaction equations involved in artificial cells. The scale bars were 20 μm.

The above cascade enzyme reactions are commonly used in this field as the model reactions for mimicking metabolism and chemical communications, which do not exist in living systems. Nicotinamide adenine dinucleotide (NADH) molecules are the products of glycometabolism, which play an important role in maintaining cell growth, differentiation, energy metabolism, and cell protection^25,26^. As one of the most important coenzymes in living cells, NADH conversion from the oxidized (NAD^+^) to reduced (NADH) forms is a common reaction in biological redox reactions. The cascade cycling of NAD^+^/NADH provides power for redox reactions, directly or indirectly regulates biochemical processes in cells and has a high correlation with the metabolic network. In living cells, the classical NADH-producing pathways involve glucose, glutamine, or fat oxidation. Herein, NADH was produced from the photosynthetic glucose inside cyanobacteria-containing artificial cells. Consequently, a NADH-dependent reaction of converting pyruvate to lactate was realized, which served as a major circulating carbohydrate fuel in mammals^27,28^. The artificial cells contained cyanobacteria, glucose dehydrogenase (GDH), and NAD^+^ loaded DPPC vesicles (1 μm in diameter), as shown in Figure 5a (left image). To avoid the NADH production during the light illumination, NAD^+^ was encapsulated inside DPPC vesicles. At room temperature, NAD^+^ was confined inside the DPPC vesicles, while at 45°C NAD^+^ were released from DPPC vesicles into cytoplasm^29^. After light illumination for 24 hours, NAD^+^ was reduced to NADH by transferring hydrogen from glucose to NAD^+^ catalyzed by glucose dehydrogenase (GDH) at 45°C for 20 minutes (Figure 5b, right image of 5a). The blue color in Figure 5b confirmed the production of NADH, which emits blue light at 460 nm. For the first time, the production of NADH was realized by using a cyanobacteria-based biological photosystem in the artificial cells. The real-time amount of produced NADH in the artificial cells (Figure 5a) was monitored using fluorescence spectrometer, which indicated a continuous increase against illumination time up to 24 hours. In the control experiments (either with no NAD^+^ or GDH), no NADH was observed (Figure 5c). The produced NADH was used to participate in the downstream metabolic reactions. The lactate dehydrogenase (LDH, 10 μg/mL) and pyruvate-LUVs (1 μm in diameter) were introduced into the artificial cells, as shown in Figure 5d (left image). NADH was used to reduce pyruvate to lactate catalyzed by LDH (right image of Figure 5d, equations in Figure 5f). The reaction was monitored by the decrease of fluorescence intensity of NADH inside the artificial cells (Figure 5e) under a fluorescence microscope. Representative fluorescence images of artificial cells (with and without pyruvate (73.1 μM)) were shown in Figure 5e. An obvious fluorescence decrease in the blue channel was observed inside the artificial cells containing pyruvate compared with the control samples (with no LDH and pyruvate-LUVs). Compared with the control samples, the fluorescence intensity of NADH was decreased by 15.85% and 18.55% at the concentration of pyruvate inside GUVs of 36.5 μM and 73.1 μM in the experimental samples respectively. The intensity of exciting laser of blue channel was kept constant for all samples. The LUVs concentration was fixed during the preparation of artificial cells. Those data confirmed the oxidation of NADH for converting pyruvate into lactate. The HPLC (high performance liquid chromatography) results of artificial cells with pyruvate (73.1 μM) showed a clear lactate peak, which provide the direct evidence for the oxidation of NADH (Figure S6). The control sample exhibited no lactate peak. The produced lactate could participate in the tricarboxylic acid cycle^30^.

**Figure 5.**
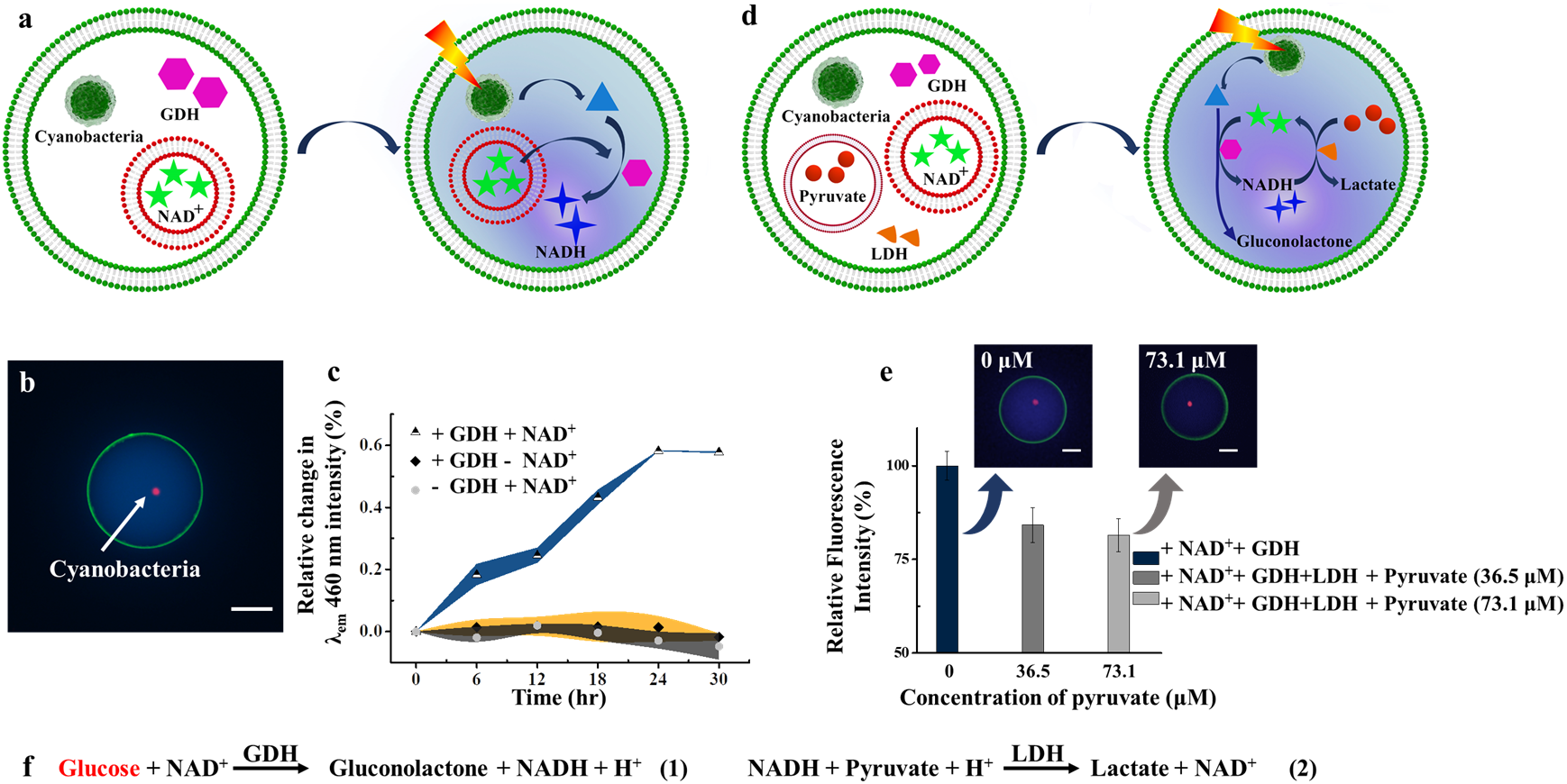
The sustainable cascade cycling of NADH/NAD^+^ inside cyanobacteria-containing artificial cells. (a) The scheme of NADH production inside an artificial cell. (b) Fluorescence microscope image of a cyanobacteria-containing artificial cell after 24 hours continuous daylight lamp illumination. Green, blue and red fluorescence represent phospholipid membrane, NADH and the cyanobacteria, respectively. (c) The normalized λ_em_ (460 nm) intensity (%) versus time curves in the artificial cells. (d) The scheme of lactate production inside an artificial cell. (e) The fluorescence microscope images of artificial cells (with and without pyruvate (73.1 μM)), and the plots of relative fluorescence intensities at different pyruvate concentration (0, 36.5 μM, 73.1 μM) in artificial cells containing cyanobacteria, NAD^+^, GDH, LDH. (f) Chemical reaction equations involved in the artificial cells. The scale bars were 20 μm.

## CONCLUSIONS

An artificial cell containing cyanobacteria was fabricated to mimic endosymbiosis. This artificial cell was able to sustainable converting light energy to chemical energy upon light illumination due to the photosynthetic capability of cyanobacteria inside the artificial cell. The “metabolism pathways” of produced glucose molecules were investigated by one enzyme cascade reaction to produce resorufin molecules, as well as one enzyme reaction to produce “cell energy currency” of NADH molecules. Furthermore, NADH was used as energy currency to convert pyruvate to lactate catalyzed by LDH. The product of NAD^+^ reentered the process of NADH production. The sustainable cascade cycling of NADH/NAD^+^ was realized, which was important for metabolic network inside cells. The developed artificial cell provides great opportunity to investigate the metabolism mechanism of cells and build sustainable energy supplied complicated minimal cells.

## Methods

### Materials

N-(7-nitrobenz-2-oxa-1,3-diazol-4-yl)-1,2-dihexadecanoyl-sn-glycero-3-phosphoethanolamine triethylammonium salt (NBD-PE) was supplied by Thermo Fisher Scientific Inc. (USA). 1,2-dimyristoyl-*sn*-glycero-3-phosphocholine (DMPC), cholesterol (Chol) and 1,2-dipalmitoyl-sn-glycero-3-phosphocholine (DPPC) were purchased from Avanti Polar Lipids, Inc. (USA). Galactose was purchased from Gentihold Biological Technology Co. Ltd. (China). Pyruvate was purchased from Aladdin Biochemical Technology Co., Ltd. (China). L-lactate dehydrogenase (LDH) was purchased from Beijing Solarbio Science & Technology Co., Ltd. (China). Glucose oxidase (GOx) from *Aspergillus niger*, horseradish peroxidase (HRP), glucose (GO) assay kit, glucose dehydrogenase (GDH), bovine serum albumin (BSA), chloroform, β-Nicotinamide adenine dinucleotide (NAD^+^), fluorescein diacetate (FDA) and Amplex red were obtained from Sigma-Aldrich, Co. (USA). Sucrose was purchased from Xilong chemical industry Co., Ltd. (China). Triton X-100 was obtained from Shanghai Haoyang Biological Technology Co. Ltd. (China). Phosphoric acid was purchased from Tianjin Fuyu Fine Chemical Co. Ltd. (China). Acetonitrile (ACN, LC-MS Grade) was supplied by Thermo Fisher Scientific Inc. (USA). Nuclepore track-etched membranes were obtained from Whatman International Ltd. (UK). Glass slides coated with indium tin oxide (ITO, sheet resistance ≈ 8 to 12 Ω per square, thickness ≈ 160 nm) were supplied by Hangzhou Yuhong Technology Co. Ltd. (China). Synechocystis sp. Strain PCC6803 was supplied by Freshwater Algae Culture Collection at the Institute of Hydrobiology (FACHB, China). Blue-Green Medium (BG-11 medium) was purchased from Qingdao Hope Bio-Technology Co., Ltd. (China). Ultrapure water from Millipore Milli-Q IQ7005 (USA) with resistivity equal to 18.2 MΩ·cm was used throughout to make solutions.

### Cultures and staining of Synechocystis sp. Strain PCC6803

Synechocystis sp. Strain PCC6803 as one type of cyanobacteria was used in this paper. Under aseptic condition, the PCC6803 cells were cultured in BG-11 medium by shaking in the sealed flask at 25°C under the daylight lamp (96 μW/cm^2^) with 12 h light on and 12 h light off in turns for 25 days. The PCC6803 cell culture solutions were then centrifugated with 4000 rpm for 5 min to remove BG-11 medium. The obtained PCC6803 cells were washed with fresh medium 3 times by centrifugation method. Consequently, the PCC6803 cells were resuspended in the mixture solution containing fresh BG-11 medium and sucrose (300 mM) (v/v = 2:8) for the preparation of artificial cells. PCC6803 cells were dyed with FDA (8 μg/mL in PBS buffer for 30 min) to indicate living cells.

### Construction of artificial cells containing cyanobacteria

Artificial cells were prepared by the electroformation method described elsewhere^31,32^. DMPC (1 mg), Chol (0.25 mg) and NBD-PE (0.05 mg) were dissolved in 200 μL chloroform. 6.5 μL such lipid mixture solution was gently spread out on the conductive surface of ITO electrodes, which were dried in a vacuum chamber for 2 h to remove chloroform. A rectangular polytetrafluoroethylene (PTFE) frame was placed between two lipid film coated ITO electrodes to form the setup. An AC electric field (4.5 V, 100 Hz) was applied by a signal generator for 2 h. Different solutions were filled inside the frame to prepare the wanted artificial cells. Four types of artificial cells were prepared. The forming solution for Type 1 contained GOx (12 μg/mL), HRP (11.88 μg/mL), cyanobacteria (22.95 mg/mL), and the mixture of sucrose (300 mM) and BG-11 medium (v/v = 8/2). The type 2 artificial cells were prepared by forming GUVs containing GOx (12 μg/mL) and cyanobacteria (22.95 mg/mL) and the mixture of sucrose (300 mM) and BG-11 medium (v/v = 8/2), followed by inward deformation of GUVs to obtain an inner vesicle containing HRP (60 μg/mL). The type 3 artificial cells contain GOx (12 μg/mL), GDH (10 μg/mL), NAD^+^-LUVs (large unilamellar vesicles), and the mixture of sucrose (300 mM) and BG-11 medium (v/v = 8/2) in the lumen. Pyruvate-LUVs and LDH (10 μg/mL) were added into type 3 artificial cells to form Type 4 artificial cells.

NAD^+^ loaded LUVs (NAD^+^-LUVs) or pyruvate loaded LUVs (pyruvate-LUVs) were prepared by an extrusion method ^33^. 1 mL DPPC and chol with a molar ratio of 7:3 (1 mg/ml) chloroform solution was dried in a vial. 1 mL of sucrose solution (300 mM) containing NAD^+^ (10 mM) or pyruvate (50 mM and 100 mM) was added to make a lipid suspension, followed by extrusion through a polycarbonate filter with a pore size of 1 µm for 21 times back and forth to obtain NAD^+^-LUVs or pyruvate-LUVs. The diameter of LUVs was 0.98 ± 0.11 μm from dynamic light scattering (DLS) measurement (Figure S7a). Unencapsulated enzymes, pyruvate and NAD^+^ were removed by centrifugation. The concentration of pyruvate (36.5 μM and 73.1 μM) in artificial cells was calculated based on the number of encapsulated pyruvate-LUVs (21.81 ± 7.67) and the size of artificial cells (24.2 ± 6.35 μm) (Figure S7b and c). A UV-Vis spectrometer was used to confirm the complete removal of enzymes and pyruvate by monitoring the characteristics absorbance of enzymes and pyruvate (Figure S8).

### Characterizations

Fluorescence images were taken under a fluorescence microscope (Olympus IX7, Japan) and confocal laser scanning microscope (Olympus FV3000, Japan). Cary 60 UV−vis spectrophotometer (Agilent, USA) and fluorescence spectrometer (PerkinElmer LS55, USA) were used for UV−vis spectra and photoluminescence spectroscopy measurements. The diameter of LUVs was obtained by Dynamic Light Scattering instrument (Malvern Zetasizer Nano, UK). The HPLC analysis was performed on a Ulimate3000 HPLC system (Thermo Fisher Scientific, USA). NIS Elements software was used to measure the fluorescence intensity of artificial cells. Video was processed by ImageJ software. All slides used for imaging were coated with BSA.

## Acknowledgements

We acknowledge the financial support provided by the National Natural Science Foundation of China (Grant No. 21773050, 21929401), and the Natural Science Foundation of Heilongjiang Province for Distinguished Young Scholars (JC2018003).

## Author contributions

X.J.H. conceived, designed and supervised the project. B.Y.Y., S.B.L. performed the experiment. B.Y.Y., S.B.L., W.M., Z.W., X.J.H. performed data analyses. B.Y.Y., X.J.H. wrote and prepared the manuscript. All authors read, revised and approved the manuscript.

