## supplementary information for "An artificial cell containing cyanobacteria for endosymbiosis mimicking"

**Movie S1**

The movie (5 frames per second, 23 seconds in real time) confirmed that the cyanobacterium was encapsulated inside the artificial cell.

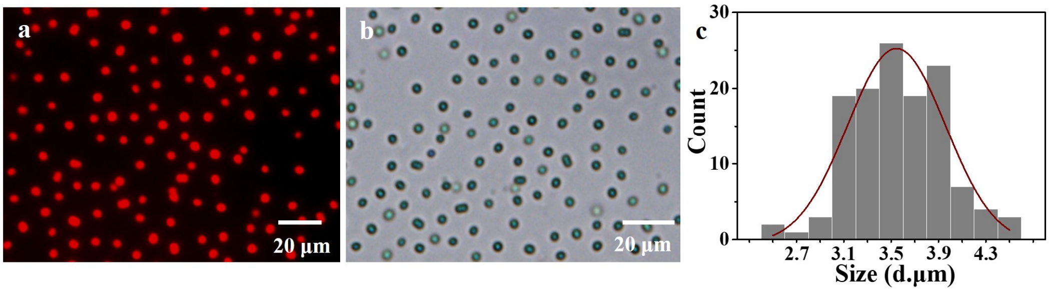


**Figure S1.** Representative fluorescence (a) and bright field (b) microscopy images of cyanobacteria. (c) Histogram of cyanobacteria cells diameters obtained from microscopy images with sample size of 127 cells.

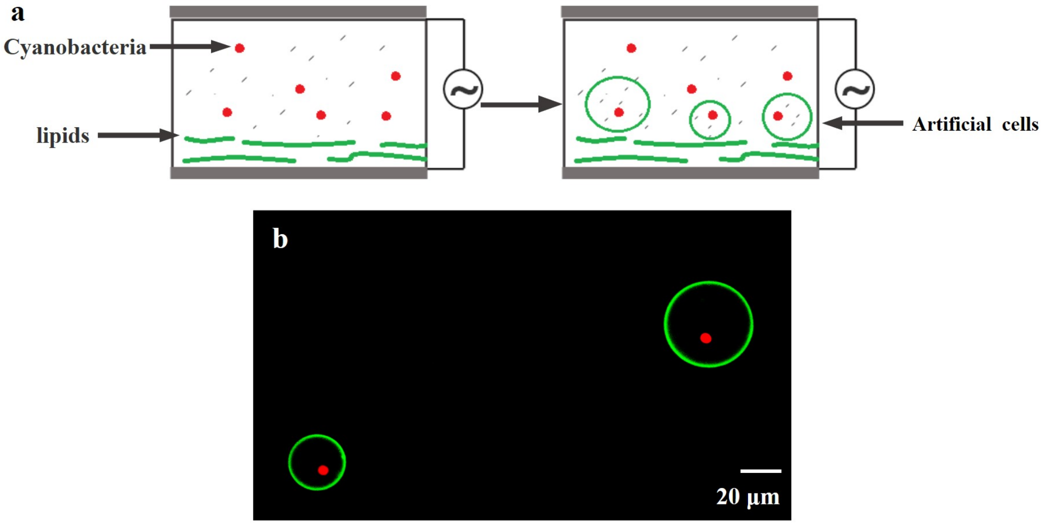


**Figure S2.** (a) Schematic illustration of the experimental setup for cyanobacteria containing artificial cells by electroformation method. (b) Representative fluorescence microscopy images of cyanobacteria containing artificial cells.

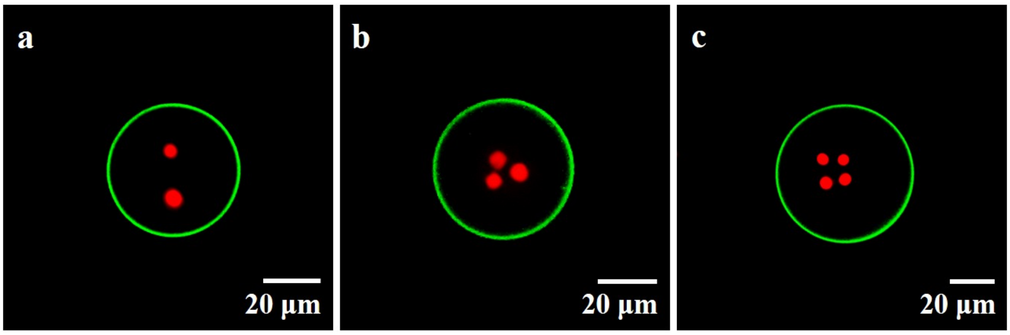


**Figure S3.** Cyanobacteria containing artificial cells. Representative fluorescence microscopy images of the artificial cells containing two (a), three (b), and four (c) cyanobacteria.

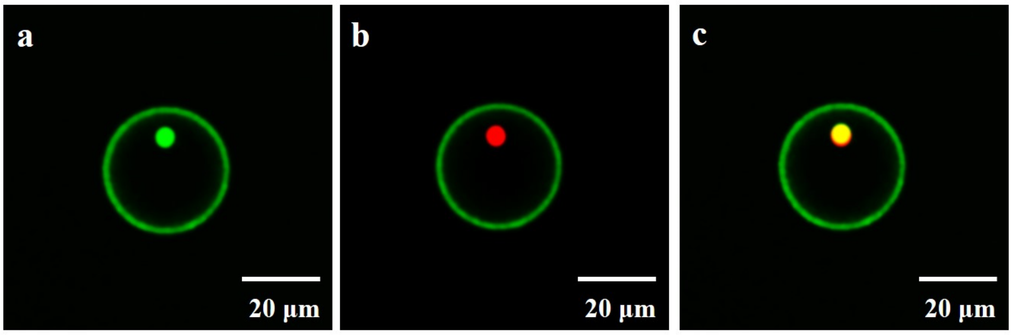


**Figure S4.** Representative artificial cell containing FDA-stained cyanobacterium after 36 hours culture. The fluorescence microscopy image with green filter (a), red filter (b), and their merged image (c). The green fluorescence of cyanobacterium indicates that it is alive.

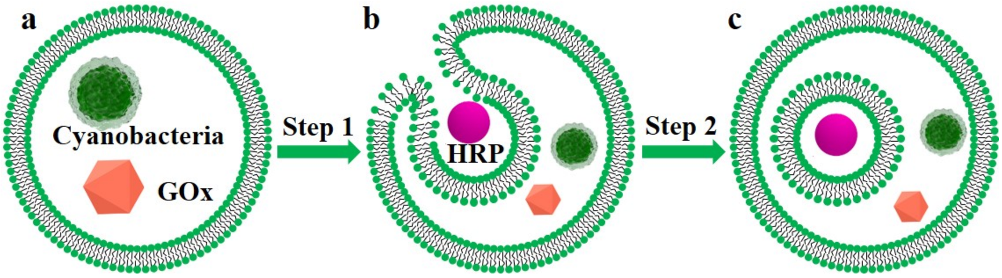


**Figure S5.** Schematic illustration of the formation of cyanobacteria artificial cells containing protoorganelle triggered by osmotic stress.

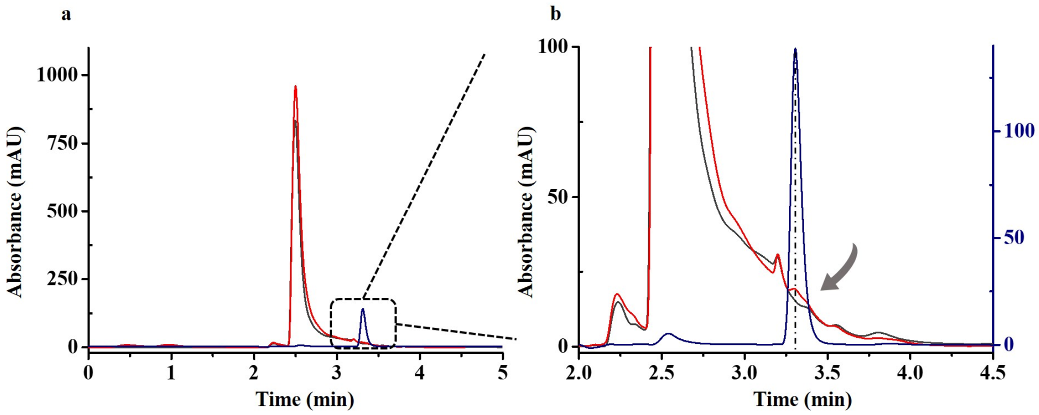


**Figure S6.** (a) HPLC chromatograms of the standards (blue), the samples with 73.1 μM pyruvate (red) and without pyruvate (black) inside artificial cells. (b) The zoom-in curves in the dotted box of (a). The clear peak at retention time of 3.330 min in the red sample curve is identical to that of the standard sample, whilst the control sample does not show any peak at this position. This result confirmed the production of lactate, and the reduction of NADH.

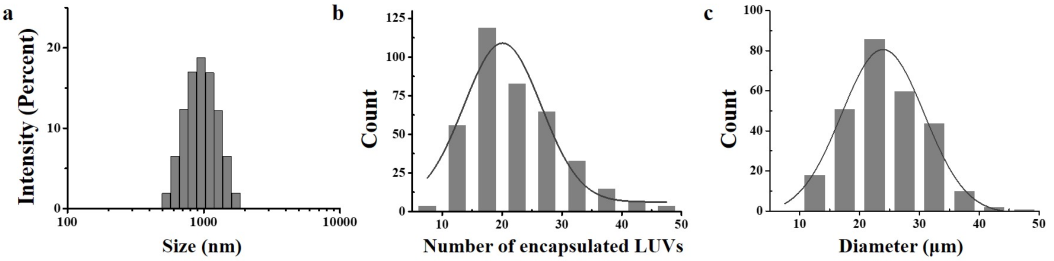


**Figure S7.** (a) Dynamic light scattering (DLS) data of LUVs. (b) Histogram of the number of encapsulated pyruvate-LUVs in artificial cells obtained from microscopy images with sample size of 386. (c) Histogram of artificial cells diameters obtained from microscopy images with sample size of 272. The mean diameter of GUVs is 24.2 ± 6.35 μm. The mean number of encapsulated LUVs inside GUVs is 21.81 ± 7.67.

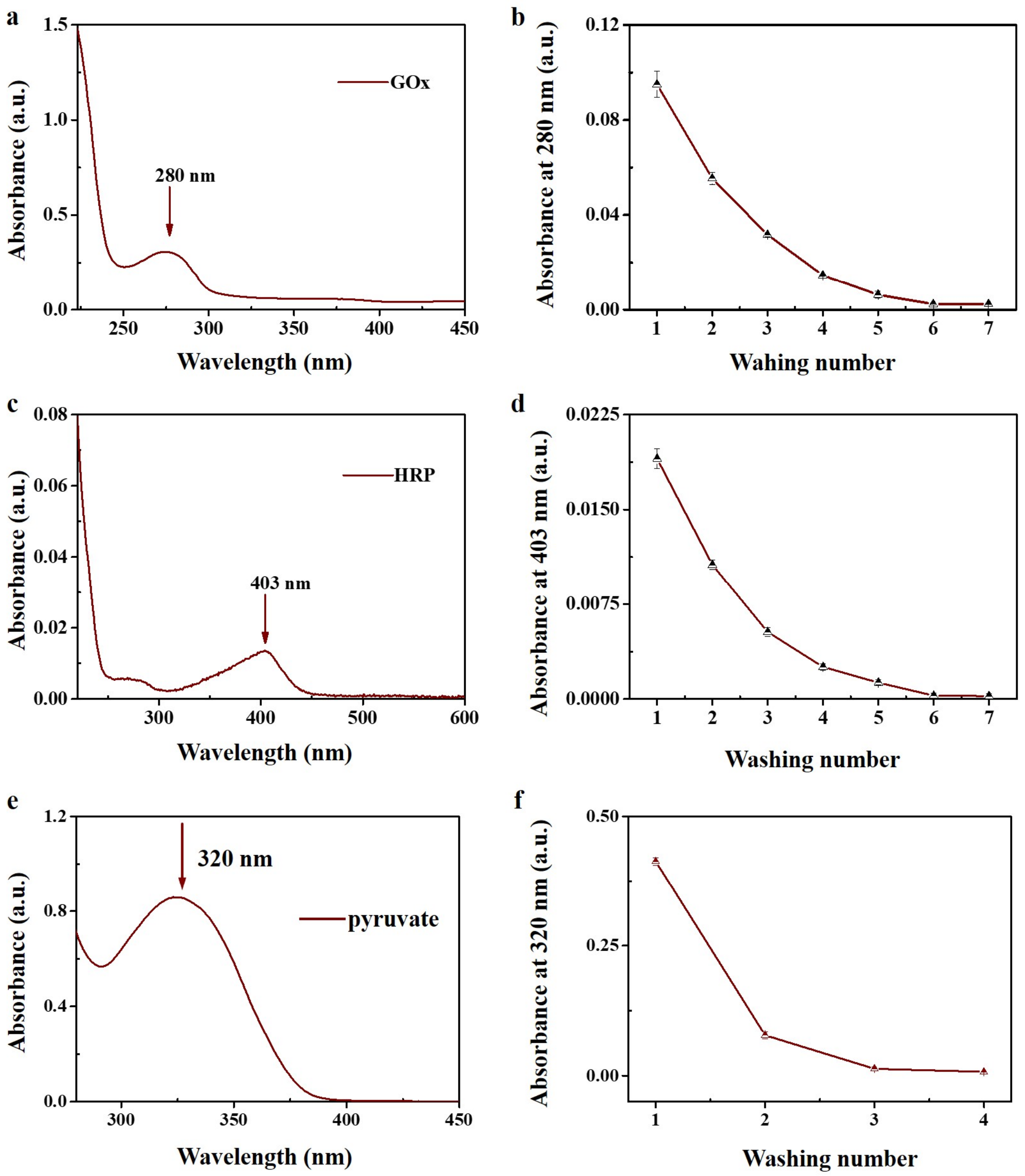


**Figure S8.** The UV-vis spectra of 18 μg/mL GOx solution (a), 11.88 μg/mL HRP solution (c) and 100 mM pyruvate solution (e). Plots of UV/Vis absorption at 280 nm for GOx (b), 403 nm for HRP (d) and 320 nm for pyruvate (f) of the supernatant after each washing by centrifugation, showing there was no residual GOx, HRP or pyruvate in the solution.
